# Intramuscular Delivery of BMP-2 and Increasing Doses of LECT-1 Using Keratin-PEG Gels for Ectopic Tissue Differentiation

**DOI:** 10.64898/2026.07.05.731787

**Authors:** Aidan Mathews, Lauren Fisher, Dana Saparova, Alejandra Cevahir, Allison Meer, Nancy Radecker, Roche C. de Guzman

## Abstract

Producing bone and cartilage in a controlled and localized manner remains a significant challenge in regenerative medicine. This study investigated the ability of keratin- and polyethylene glycol (PEG)-based degradable hydrogels to deliver bone morphogenetic protein 2 (BMP-2) and leukocyte cell-derived chemotaxin 1 (LECT-1; also known as chondromodulin-1) intramuscularly to induce ectopic tissue formation. Adult male CD-1 mice received intramuscular implants of keratin-PEG gels containing a fixed dose of BMP-2 and increasing amounts of LECT-1. After two weeks, implants and surrounding muscle were analyzed using computed tomography (CT) and histology. The results showed that BMP-2 is necessary for forming new bone and cartilage, whereas LECT-1 alone appeared to trigger muscle dedifferentiation without ossification or chondrogenesis. Co-delivery of BMP-2 and LECT-1 enhanced bone and cartilage formation in a dose-dependent manner: higher LECT-1 doses led to proportionally more ectopic cartilage (linear correlation, r^2^ ≈ 90%), while bone formation peaked at the third LECT-1 dose at approximately twice the volume of the BMP-2-only group. These findings indicate that muscle-resident cells may be capable of reverting and switching to mesenchymal lineages, recapitulating endochondral ossification. The platform offers a promising strategy for growing bone and cartilage autografts within skeletal muscle bundles.

**Impact Statement:** This study presents a novel strategy for inducing ectopic bone and cartilage formation by delivering BMP-2 and LECT-1 from keratin-PEG hydrogels into skeletal muscle. The findings suggest that muscle-resident cells can dedifferentiate and transdifferentiate into mesenchymal lineages, enabling controlled ectopic tissue regeneration. This approach may inform future tissue engineering therapies in which autografts are harvested from within the body, using skeletal muscle as an *in vivo* tissue incubator.

## Introduction

Bone fractures and musculoskeletal injuries represent a major global health burden, with an estimated 178 million new fractures occurring annually worldwide.^1^ Accelerating and improving bone and cartilage regeneration remains a critical goal in regenerative medicine and tissue engineering. Bone formation occurs through two primary developmental pathways: intramembranous ossification, which converts mesenchymal tissue directly into bone, and endochondral ossification, which proceeds through an intermediate cartilage template that is later replaced by bone.^2^ Endochondral ossification is particularly relevant for long-bone development and fracture healing, where a cartilage anlage precedes ossification, and it has increasingly been exploited as a developmental engineering route for the repair of large bone defects.^3^

The cellular contributors to these processes include mesenchymal stem cells (MSCs), osteogenic progenitors, osteoblasts, osteoclasts, and osteocytes. MSCs are multipotent and can differentiate into osteoblasts, chondrocytes, adipocytes, and myoblasts depending on local signaling cues.^4^ Skeletal muscle contains satellite cells, the quiescent muscle-resident stem cells that activate upon injury and contribute to regeneration.^5^ Beyond their myogenic role, satellite cells and muscle-derived progenitors have been shown to be multipotential, adopting osteogenic, chondrogenic, and adipogenic fates under appropriate stimuli.^6^ These cells, along with interstitial fibro-adipogenic progenitors (FAPs) and other muscle-resident mesenchymal populations, can influence the fate of implanted biomaterials and associated growth factors.^7-9^ Cartilage integrity depends on chondrocytes, which synthesize extracellular matrix (ECM) components such as type II collagen and proteoglycans. Degenerative joint diseases such as osteoarthritis result from ECM degradation, emphasizing the need for strategies that promote robust cartilage formation.^10^

Growth factor-based therapies have emerged as promising approaches to direct stem cell differentiation and enhance tissue regeneration. Bone morphogenetic protein 2 (BMP-2) is a well-characterized osteoinductive factor used clinically to stimulate bone formation.^11^ However, the burst release of BMP-2 from its standard absorbable collagen sponge carrier has been associated with off-target effects, including soft-tissue swelling, inflammation, and uncontrolled heterotopic ossification, motivating the search for carriers that provide localized and sustained delivery.^12,13^ In contrast, leukocyte cell-derived chemotaxin 1 (LECT-1), more commonly known as chondromodulin-1, is less studied but exhibits chondrogenic potential together with anti-angiogenic properties, maintaining the cartilage phenotype by inhibiting vascular invasion.^14-16^ Despite their individual roles in osteogenesis and chondrogenesis, the combined effects of BMP-2 and LECT-1 in ectopic environments remain poorly understood. Intramuscular implantation models provide a unique platform to study these interactions, allowing evaluation of tissue formation in a non-skeletal, vascularized environment.

Keratin extracted from human hair is a biocompatible, tunable, and bioactive protein biomaterial that has been used for drug delivery, wound healing, and bone regeneration, and that binds and sequesters BMP-2 to control its release.^17-22^ Combining keratin with photopolymerized polyethylene glycol (PEG) yields a degradable matrix that can provide localized and sustained growth-factor presentation, analogous to degradable PEG hydrogels previously shown to deliver osteoinductive factors and drive ectopic mineralization.^23^

The objective of this study was to investigate the tissue differentiation responses to intramuscular co-delivery of BMP-2 and increasing doses of LECT-1 using keratin-PEG gels. We hypothesized that LECT-1 would promote cartilage formation in a dose-dependent manner and that its combination with BMP-2 would recapitulate endochondral ossification, leading to mature bone formation via a cartilage intermediate. Additionally, we explored whether the anti-angiogenic properties of LECT-1 could modulate local tissue remodeling and enhance the biocompatibility of the hydrogel construct.

## Materials and Methods

### Gel implants with growth factors

Keratin-PEG gels (G) were prepared using a modified thiol-ene photopolymerization protocol.^17,18^ Briefly, reduced keratin-based extract (kerateine) was obtained from human hair fibers following thioglycolic acid reduction, dialysis, and concentration, yielding soluble protein extracts with an average molecular weight (M_w_) of ∼98 kDa and an isoelectric point (pI) of 5.3. Polyethylene glycol diacrylate (PEGDA, M_w_ = 700 Da) was used as the synthetic precursor. The pregel solution contained 10 mg/mL kerateine, 20% (V/V) PEGDA, 5% (V/V) glycerol, and 2% (m/V) Irgacure 2959 photoinitiator (from a 10% stock in methanol) in 10 mM NaOH. Growth factors (GFs) were added prior to crosslinking: BMP-2 (first growth factor, GF1: B; Infuse system, Medtronic, Minneapolis, MN) at a fixed dose of 25 µg, and/or LECT-1 (MyBioSource, San Diego, CA) at increasing doses of 25, 50, 100, and 200 ng per implant (second growth factor, GF2: L1 [lowest], L2, L3, and L4 [highest], respectively). Solutions were filter-sterilized (0.22-µm filter), dispensed into sterile 24-well plates (750 µL per well), and exposed to UV light (365 nm, 20 min) to induce PEG chain growth and thiol-ene crosslinking with keratin. Gels were stored in a humidified environment at 4°C until use. On the day of implantation, cylindrical gels (5 mm diameter × 4 mm height; ∼77 µL volume and ∼101 mm^2^ surface area) were obtained using sterile biopsy punches and equilibrated to room temperature.

### Animal study on intramuscular implantation

All procedures were approved by the Institutional Animal Care and Use Committee (IACUC). Adult male CD-1 IGS albino mice (Charles River, Wilmington, MA; 12 weeks old, 40 ± 3 g) were anesthetized with isoflurane (4% induction, 1% maintenance) using a SomnoSuite low-flow system (Kent Scientific, Torrington, CT). After hair clipping and povidone-iodine disinfection, a 15-mm incision was made over the left hamstring (posterior femoral muscles) region. A blunt probe was used to separate the fascia of the biceps femoris and semitendinosus muscles, creating a 10-mm intramuscular (IM) pocket without damaging muscle fibers. One gel implant (**Table 1**) was inserted per pocket. The separated muscles were drawn back together and closed with degradable 4-0 polyglactin sutures (two separate knots) to keep the gel in place, and the skin was closed with interrupted nylon sutures. Postoperative care included monitoring and analgesia (subcutaneous 5 mg/kg ketoprofen for 3 days). Implants remained *in situ* for 14 days. At the two-week endpoint, the animals were euthanized. The left leg with the implant, surrounding muscles, and bone was collected, fixed in 10% neutral-buffered formalin (Sigma-Aldrich, St. Louis, MO) for 7 days, washed in phosphate-buffered saline (PBS), and stored in 60% (V/V) ethanol.

**TABLE 1.**
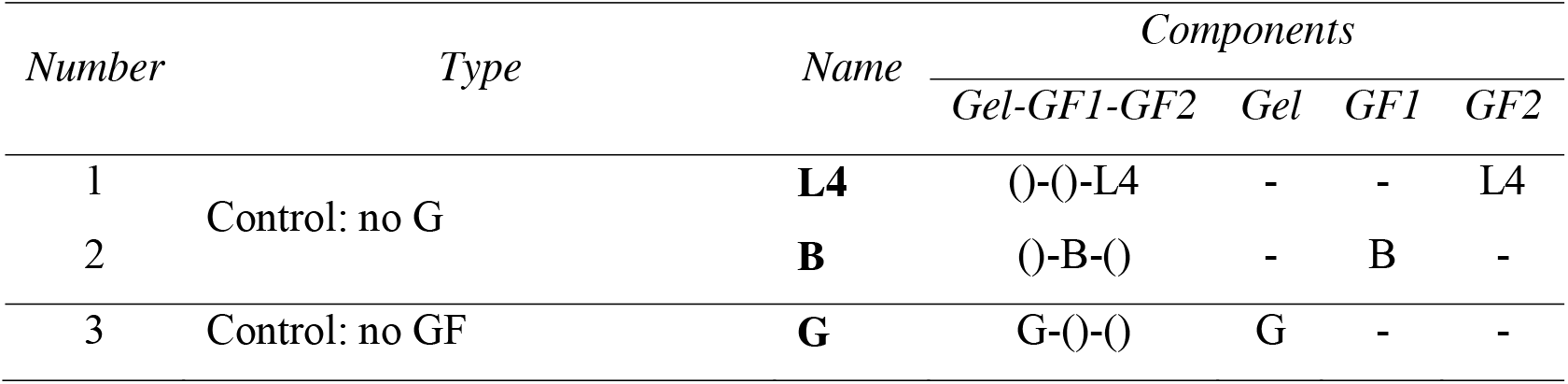
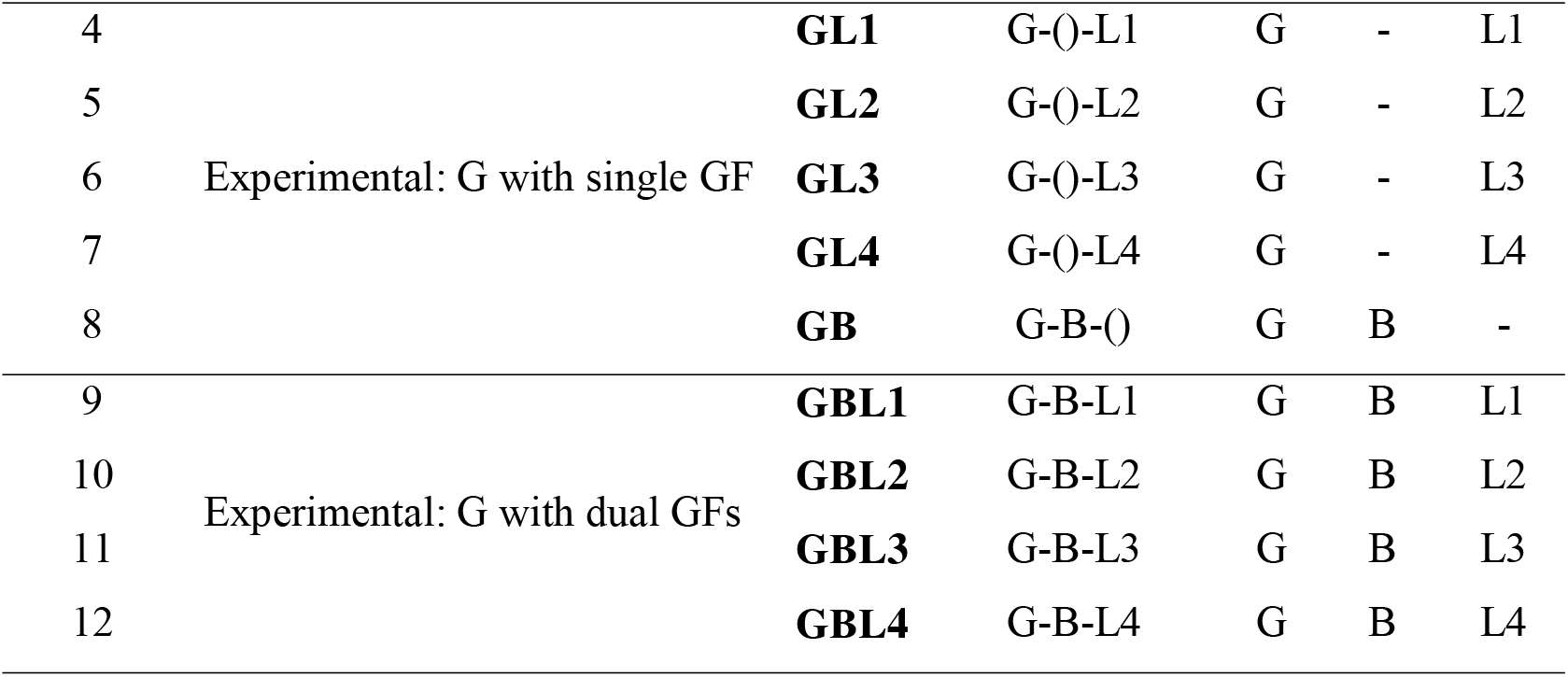
Animal treatment groups. Dashes (-) indicate an absent component.

### Control and experimental groups

Twelve animal treatment groups (**Table 1**) were tested at n = 4 mice (biological replicates) per group: *no-gel controls* (77 µL of liquid solution injected into the muscle pocket) at the highest dose of LECT-1 (L4, 200 ng) and BMP-2 only (B, 25 µg); a *no-growth-factor control* (keratin-PEG gel only, G); *gel with single growth factor experimental groups* (G with increasing doses of LECT-1, GL1–GL4, and G with BMP-2 only, GB); and *gel with dual growth factor experimental groups* (G plus BMP-2 with increasing LECT-1, GBL1 to GBL4).

### Computed tomography for bone detection

Two-week gel implants within the muscle pocket were wrapped in PBS-soaked gauze, arranged on a holder, and scanned using a micro-computed tomography (µ-CT) system (Inveon, Siemens Healthineers, Erlangen, Germany; with COBRA cone-beam reconstruction, Exxim, Pleasanton, CA) at 97.13-µm isotropic voxel resolution. DICOM (digital imaging and communications in medicine) files were analyzed in MATLAB (MathWorks, Natick, MA) to assess bone formation by measuring calcified-tissue volume and density using region-of-interest integrals after background subtraction.

### Histology of ectopic tissue formation

Histological preparation followed our previously established protocols.^19^ Specimens were trimmed to retain only the implants and adjacent muscle tissue. Decalcification was achieved using 12% (V/V) formic acid, followed by neutralization in 0.1% (m/V) sodium bicarbonate and washing with water. Samples were embedded in paraffin (Paraplast X-TRA, Sigma-Aldrich), sectioned at 5 µm using a microtome, and mounted onto glass slides. For general connective and skeletal-muscle tissue assessment, including cartilage and bone, sections were stained with Masson’s trichrome (MT), yielding black/dark staining for cell nuclei, red for cellular cytoskeleton and cytosol, and blue for collagenous ECM. To specifically identify cartilage, sections were stained with 0.05% (m/V) toluidine blue (TB) in McIlvaine buffer (disodium phosphate and citric acid) adjusted to pH 3.5, where proteoglycans and glycosaminoglycans (GAGs) appear purple (metachromatic).^24-26^ Image capture was performed using the Cytation 5 Cell Imaging Multimode Reader (Agilent, Santa Clara, CA), Primostar 3 (Zeiss, White Plains, NY), and FV3000 (Olympus, Evident Scientific, Waltham, MA).

### Data presentation and statistical methods

Image processing, analysis, and presentation were performed with GIMP (gimp.org), ImageJ (NIH, Bethesda, MD), MATLAB, and Office Excel and PowerPoint (Microsoft, Redmond, WA). For each biological sample, three technical replicates were analyzed, and results are shown as mean (AVE) ± 1 standard deviation (STD). Semi-quantitative scoring of new tissues (altered muscle, cartilage, and bone) used an ordinal scale (0 = absent; 1 and increasing integers proportional to the relative amount). Statistical comparisons used the Student’s t-test and analysis of variance (ANOVA) with Tukey-Kramer post hoc multiple-comparison tests. Significance was indicated as *p < 5%, **p < 1%, and ***p < 0.1%.

## Results

### Gel characteristics

The keratin-PEG gels appeared similar regardless of growth-factor loading. They were white-brown and translucent, mechanically stiff, and readily handleable, retaining the cylindrical geometry of the biopsy punch (5 mm diameter × 4 mm height; ∼77 mm^3^ volume) (**Fig. 1**). The photopolymerized matrix remained intact during implantation and was identifiable at the two-week endpoint as brownish remnants within the muscle pocket (**Fig. 2, arrows**).

**FIG. 1.**
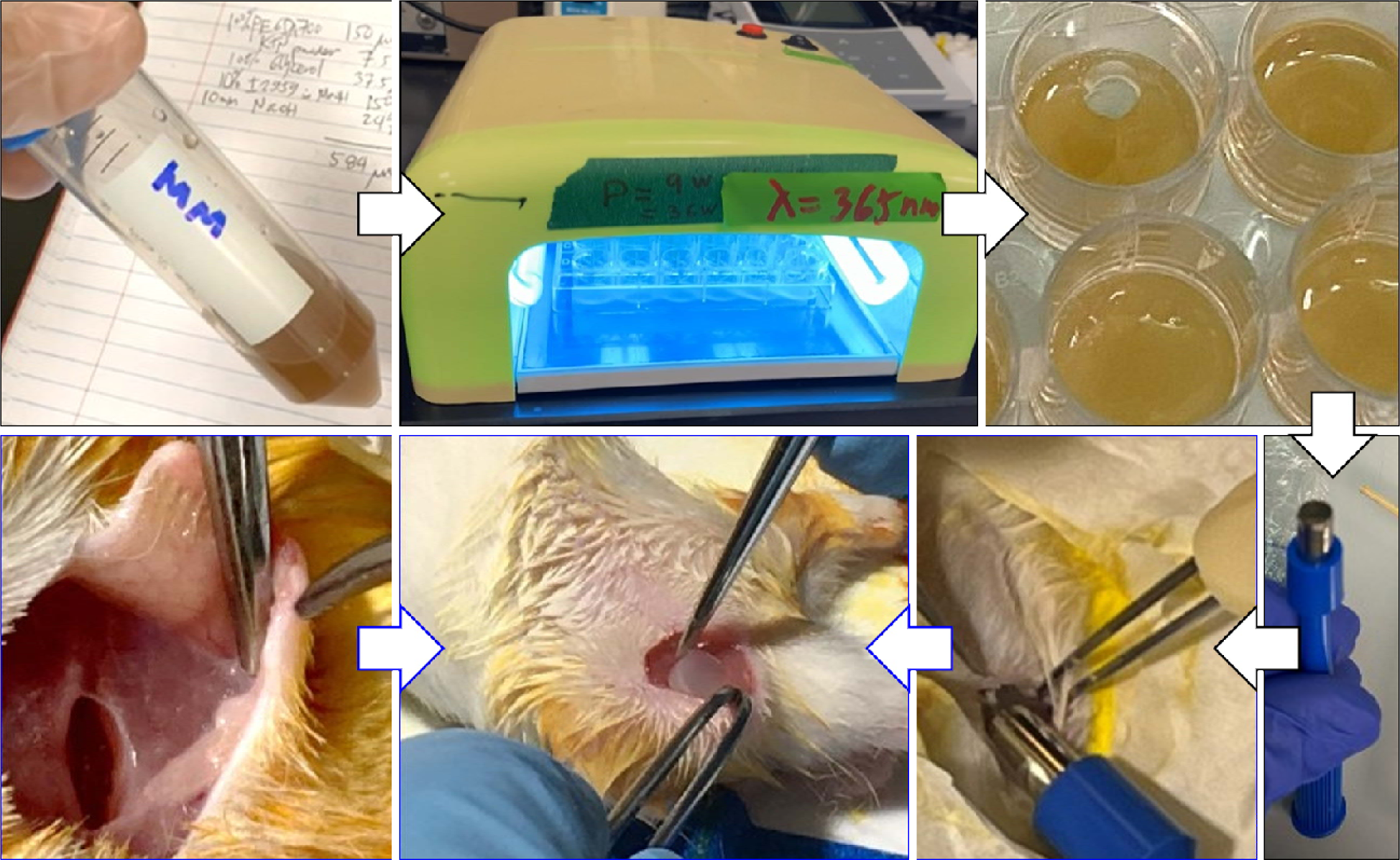
Gelation and intramuscular implantation of gel constructs. The liquid master mix (MM) was dispensed into the wells of a 24-well microplate and exposed to 365-nm light, resulting in gelation. Biopsy punches of 5-mm diameter samples were then implanted into pockets created by separating the hamstring muscles of mice.

**FIG. 2.**
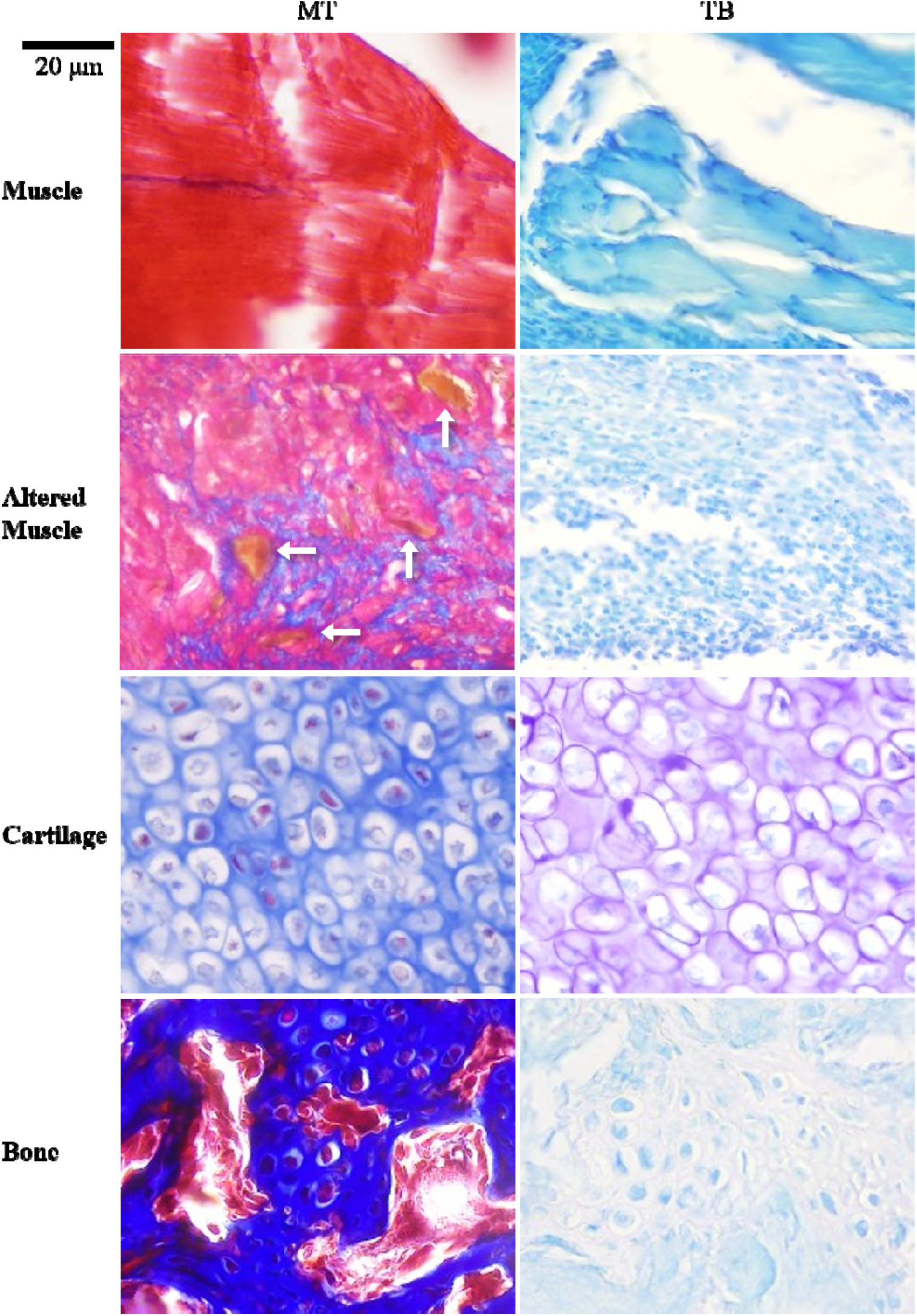
Representative histology of tissues formed in the intramuscular space, stained with Masson’s trichrome (MT; left column) and toluidine blue (TB; right column). Rows show normal muscle, altered (dedifferentiating, unbundled) muscle, neo-ectopic cartilage, and neo-ectopic bone. In MT, nuclei stain dark, cytoplasm and cytoskeleton red, and collagenous extracellular matrix blue; in TB, proteoglycan- and glycosaminoglycan-rich cartilage is metachromatic (purple). The neo-cartilage shows entrapped chondrocytes within a proteoglycan-rich matrix, and the neo-bone shows trabeculae with marrow-like luminal spaces. Arrows in the altered-muscle (MT) panel indicate brownish remnants of the gel implant at the 2-week endpoint. Scale bar = 20 µm.

### Neo-tissue morphological analysis (histology)

Histological evaluation revealed distinct tissue-differentiation patterns across the experimental groups, visualized using Masson’s trichrome and toluidine blue staining (**Fig. 2**) and scored semi-quantitatively across all twelve groups (**Fig. 3**). MT staining demonstrated collagen-rich ECM in groups treated with BMP-2, with or without LECT-1. In BMP-2-containing groups, dense blue-stained regions indicative of bone matrix were observed, together with red-stained cytoplasmic regions and dark nuclei corresponding to active osteoblasts. The neo-ectopic bone was trabecular, with marrow-like spaces in the lumen. Toluidine blue staining confirmed proteoglycan-rich cartilage, particularly in groups receiving BMP-2 in combination with increasing doses of LECT-1; the neo-ectopic cartilage contained proliferating chondroblasts and entrapped chondrocytes within a metachromatic, proteoglycan-rich matrix and was devoid of blood vessels (**Fig. 2**).

**FIG. 3.**
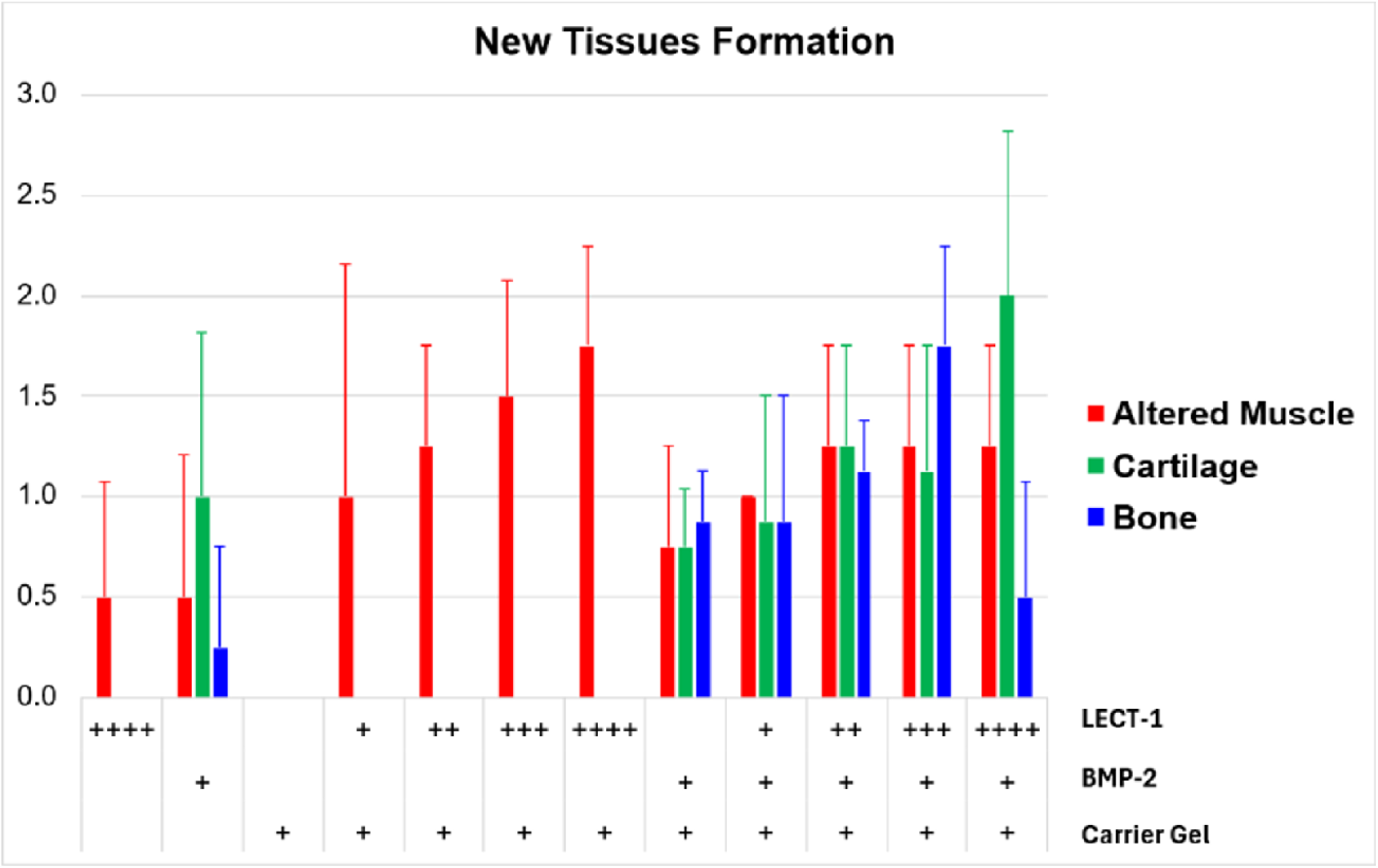
Semi-quantitative analysis of new tissue formation (altered muscle, cartilage, and bone) across the twelve treatment groups, scored on an ordinal scale (mean ± 1 standard deviation; n = 4). Groups are defined by the presence and dose of carrier gel, BMP-2, and LECT-1, as indicated beneath the axis. Altered muscle appeared across most growth-factor groups, whereas cartilage and bone arose only when BMP-2 was present; cartilage increased and bone shifted with increasing LECT-1 dose in the dual-growth-factor (gel + BMP-2 + LECT-1) groups.

In groups treated with BMP-2 alone (GB), both cartilage and bone were evident, indicating that BMP-2 is sufficient to induce ectopic ossification in muscle. The addition of LECT-1 enhanced cartilage formation in a dose-dependent manner. Increasing LECT-1 at molar ratios of 0, 1, 2, 4, and 8 relative to BMP-2 produced a linear increase in cartilage (r^2^ ≈ 90%; **Fig. 4A**), with significantly more cartilage in the highest dual-dose group than in the BMP-2-only group (*p = 0.03; **Fig. 4A**). These avascular cartilage islands were consistent with the known anti-angiogenic properties of LECT-1. Formed cartilage was located peripheral to the degraded gel or, when bone was present, in the inner region.

**FIG. 4.**
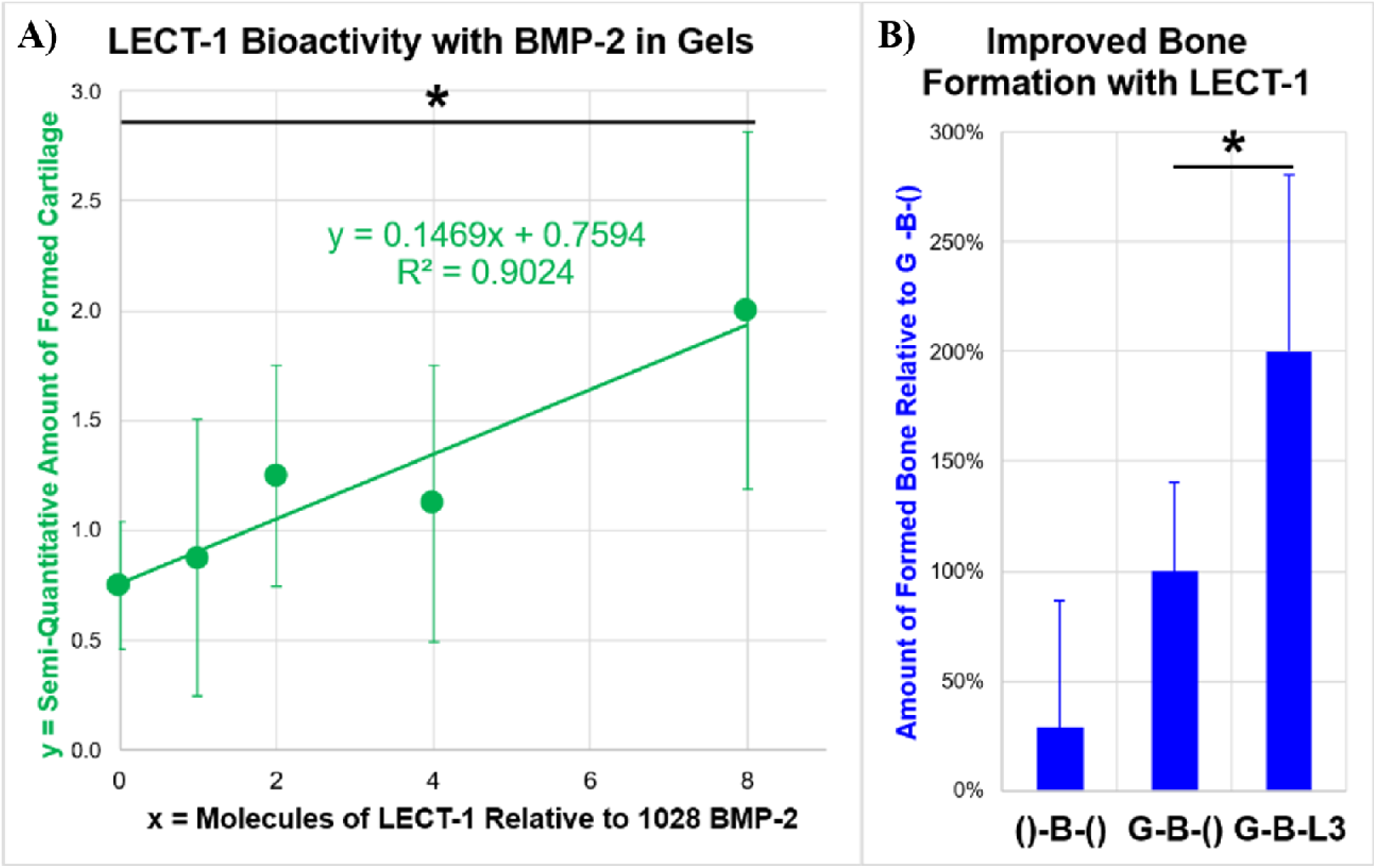
Dose-dependent effect of LECT-1 co-delivered with a fixed dose of BMP-2 in keratin-PEG gels. A) LECT-1 bioactivity: the semi-quantitative amount of formed cartilage increased linearly with the number of LECT-1 molecules relative to 1028 BMP-2 molecules (y = 0.1469x + 0.7594, r^2^ = 0.9024); cartilage was significantly greater at the highest LECT-1 dose than without LECT-1 (*p = 0.03). B) Improved bone formation: the amount of formed bone relative to the gel-with-BMP-2-only group (G-B-(), set to 100%) was ∼29% without a carrier gel and ∼200% for the dual-growth-factor group G-B-L3 (GBL3), which was significantly greater than G-B-() (*p = 0.02).

In contrast, LECT-1 delivered without BMP-2 (GL1–GL4) did not induce cartilage or bone but consistently produced “altered” muscle, characterized by unbundling of myofibers and their separation into individual cells. The extent of this dedifferentiation was dose-dependent and was most apparent at the lower LECT-1 doses. No new cartilage or bone was observed in any group lacking BMP-2, including the gel-only control (G), which showed intact implants surrounded by normal muscle.

### CT of bone features

CT imaging corroborated the histological findings, revealing mineralized tissue only in BMP-2-treated groups; no calcified tissue was detected in the controls or in groups treated with LECT-1 alone (**Fig. 5**). BMP-2-only implants showed moderate bone density, whereas BMP-2 + LECT-1 groups exhibited denser, cortical-like bone (**Fig. 5**). Without a carrier gel, BMP-2 still induced ectopic bone, but the bone was diffuse, lacked a defined shape, and formed in various locations, including adjacent to the femur (heterotopic and uncontrolled); the no-gel control contained approximately 29% of the bone volume measured in the corresponding gel implant (**Fig. 4B**). Delivery of BMP-2 from the gel produced a defined, controlled bone structure that followed the shape of the implant. Bone formation increased with LECT-1 dose up to the third dose: the dual-growth-factor group GBL3 produced approximately twice (200%) the bone of the BMP-2-only gel group GB (*p = 0.02; **Fig. 4B**). The highest LECT-1 dose (GBL4), however, yielded significantly less bone than GBL3 but markedly more cartilage, indicating that the LECT-1 dose is critical to controlling whether bone or cartilage predominates.

**FIG. 5.**
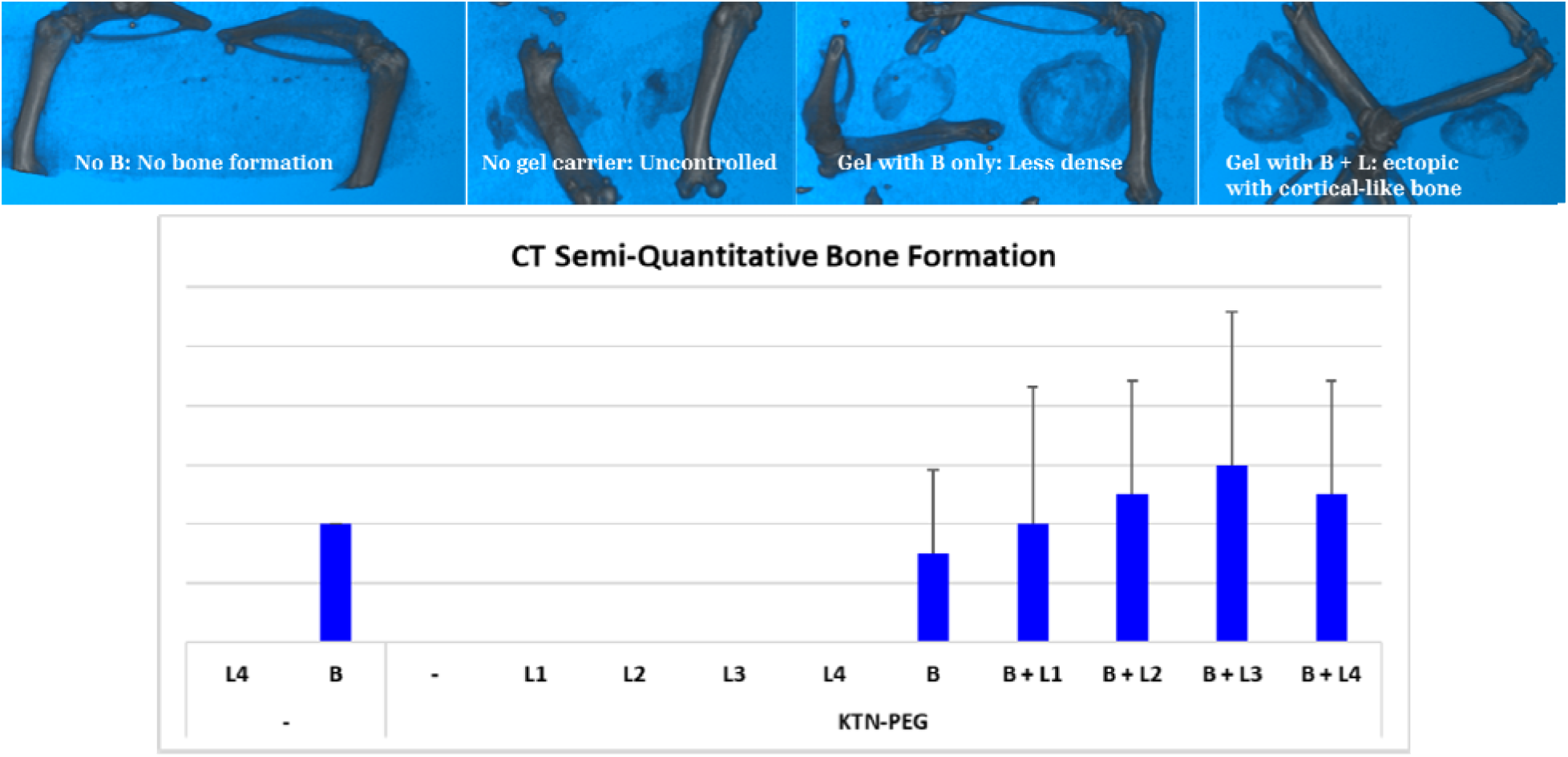
Micro-computed tomography (µ-CT) of ectopic bone. Top: representative reconstructed µ-CT images showing no mineralized tissue without BMP-2 (No B), diffuse and uncontrolled heterotopic bone when BMP-2 was delivered without a carrier gel (No gel carrier), defined but less-dense bone from the gel with BMP-2 only (Gel with B), and dense, cortical-like ectopic bone from the gel with BMP-2 + LECT-1 (Gel with B + L). Bottom: semi-quantitative CT bone formation across all groups (mean ± 1 standard deviation; n = 4); calcified tissue was detected only in BMP-2-containing groups and increased with LECT-1 dose, peaking at the third dose (B + L3).

### Summary of key observations

- BMP-2 is essential for ectopic bone and cartilage formation in skeletal muscle.
- LECT-1 alone does not induce ossification but promotes muscle dedifferentiation, potentially activating endogenous progenitor populations.
- Increasing LECT-1 doses enhance cartilage formation linearly and modulate bone density when co-delivered with BMP-2, with bone peaking at the third dose and cartilage greatest at the highest dose.
- Histological and imaging data suggest that the BMP-2 + LECT-1 combination recapitulates endochondral ossification, with cartilage serving as a precursor to bone.
- Muscle-derived satellite cells and fibroblast-like cells may serve as a source of mesenchymal progenitors in this model.

## Discussion

This study demonstrates that intramuscular delivery of BMP-2 and LECT-1 from keratin-PEG hydrogels can induce ectopic cartilage and bone in skeletal muscle, with distinct dose-dependent effects and spatial organization. The findings suggest a biologically orchestrated sequence of tissue transformation, initiated by growth-factor release and mediated by local cellular responses.

Our data confirm that the gel carrier is essential for localized and sustained delivery. In the absence of the keratin-PEG matrix, BMP-2 and LECT-1 produced diffuse and uncontrolled responses. BMP-2 led to heterotopic bone adjacent to native skeletal structures, while LECT-1 caused widespread muscle dedifferentiation. This mirrors the clinical experience with burst release of BMP-2 from collagen sponges, which has been linked to off-target ossification and soft-tissue complications,^12,13^ and supports the use of binding matrices such as keratin and degradable PEG to constrain growth-factor activity in space and time.^19,23^ Gel-only controls showed no tissue remodeling, indicating that the matrix itself is inert and does not induce ectopic tissue formation.

LECT-1 alone did not induce cartilage or bone but consistently triggered myofiber unbundling and dedifferentiation. Histological and confocal imaging revealed separation of myofibers into individual cells, activation of satellite cells, and emergence of fibroblast-like morphologies. Skeletal muscle harbors several populations capable of mesenchymal differentiation, including multipotential satellite cells^6^ and interstitial fibro-adipogenic progenitors,^7-9^ any of which could be recruited under the local signaling established by the implant. These observations suggest that LECT-1 may initiate a transdifferentiation process that converts muscle-resident cells into mesenchymal-like progenitors; the dose-dependence of the effect implies that LECT-1 modulates progenitor plasticity, possibly by altering the local microenvironment or by suppressing angiogenesis through its anti-angiogenic activity.^15,16^

BMP-2 was necessary for the induction of both cartilage and bone. BMP-2 is a potent inducer of osteogenic and chondrogenic differentiation of mesenchymal and myogenic cells, and its introduction into muscle classically produces ectopic endochondral bone.^11,27^ When delivered alone, BMP-2 produced moderate bone and cartilage. When combined with increasing doses of LECT-1, cartilage formation was enhanced and bone density increased, peaking at the third LECT-1 dose. This supports the hypothesis that LECT-1 primes the tissue environment for chondrogenesis, which then serves as a template for BMP-2-driven endochondral ossification. The spatial arrangement of tissues: cartilage in the inner region and bone on the outer surface of the degraded gel, and the continuity between them are consistent with cartilage maturing into bone, as occurs during developmental and fracture-associated endochondral ossification.^2,3,28^

BMP signaling also influences the balance between proliferation and differentiation of muscle satellite cell descendants, and can drive myogenic cells toward osteogenic and chondrogenic fates,^27,29^ providing a plausible mechanistic basis for the muscle-derived origin of the new tissues observed here. The origin of the newly formed tissues remains a key question; based on histological evidence and prior work, three potential sources are proposed: 1) muscle-resident satellite cells, which can activate and undergo lineage reprogramming;^5,6^ 2) interstitial fibroblast/FAP-type progenitors, which respond to growth-factor signaling and have demonstrated osteogenic and chondrogenic capacity;^7,8,30^ and, less likely, 3) circulating MSCs, whose contribution is constrained by the avascular nature of the cartilage regions. Lineage-tracing studies of BMP-driven heterotopic ossification have variously implicated muscle-resident and perivascular/endothelial-associated progenitors,^31,32^ underscoring that the cellular source is context-dependent. The consistent observation of muscle dedifferentiation preceding cartilage and bone formation in our model suggests that the muscle itself is a primary source of progenitor cells.

The proposed mechanism (**Fig. 6**) begins with the release of LECT-1 and BMP-2 from the gel surface, followed by muscle dedifferentiation and activation of local progenitors. LECT-1 promotes chondrogenesis and inhibits angiogenesis, leading to the formation of avascular cartilage, while BMP-2 drives ossification through endochondral pathways, resulting in mature bone. This sequence is supported by the observed histological order from the gel surface (altered muscle, bone, cartilage, gel remnants) and by CT imaging showing cartilage preceding bone and increasing tissue complexity with dual growth-factor delivery. Pharmacologic studies showing that retinoic acid receptor-γ agonists and BMP type I receptor inhibitors block ectopic chondrogenesis and ossification further support an endochondral mechanism that could, in principle, be tuned in this platform.^31,33^

**FIG. 6.**
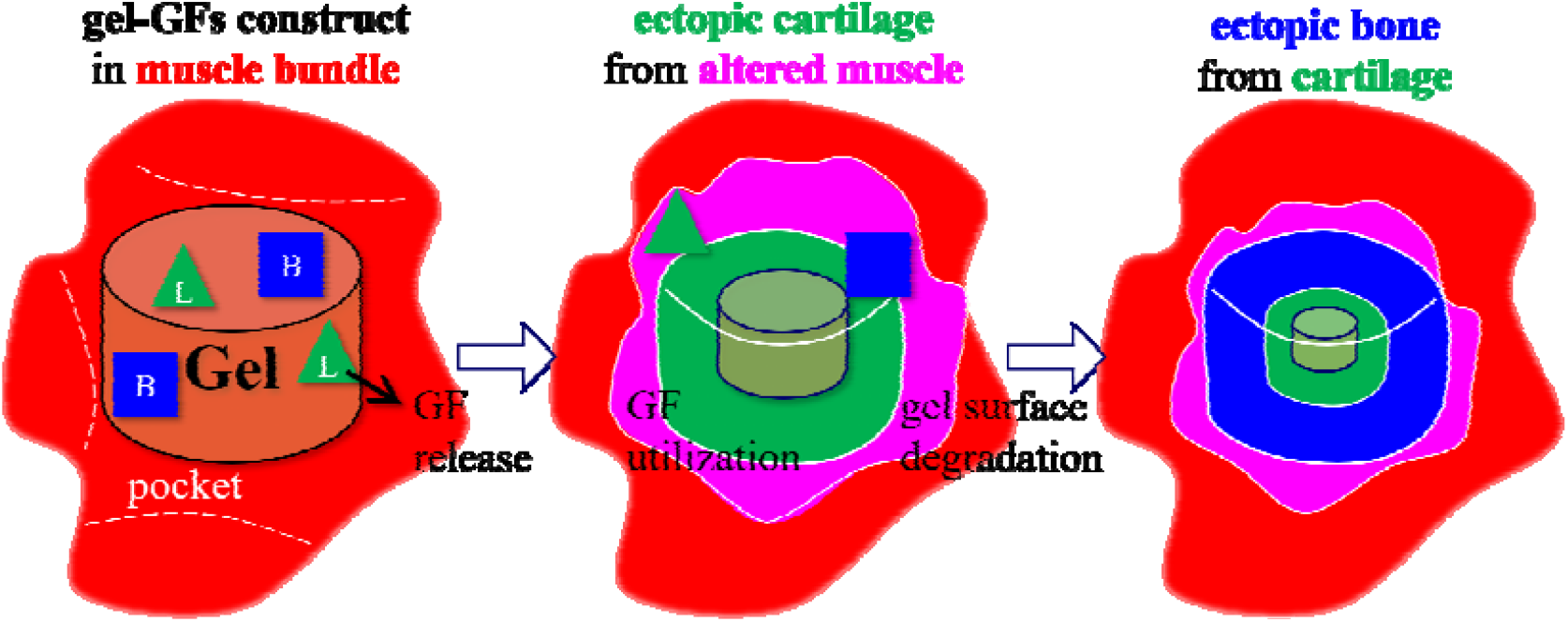
Proposed mechanism of dual growth factor (B = BMP-2 and L = LECT-1) release leading to temporal formation of altered muscle tissue first, then ectopic cartilage and ectopic bone within the skeletal muscle.

In summary, muscle tissue exposed to specific growth-factor combinations can undergo dedifferentiation and apparent transdifferentiation into cartilage and bone. The keratin-PEG gel enables spatial control of growth-factor release, and the dual delivery of BMP-2 and LECT-1 recapitulates developmental ossification pathways. A limitation of the present work is the two-week endpoint and semi-quantitative scoring; longer time points, lineage tracing, and lower-BMP-2 formulations intended to produce an exclusive cartilage response are warranted. These findings have implications for the design of biomaterials that harness endogenous cell populations for tissue repair and for the harvesting of ectopic autografts to treat critical-size defects.

## Conclusion

Keratin-PEG hydrogels provide a controllable, biocompatible platform for the localized intramuscular co-delivery of BMP-2 and LECT-1. BMP-2 was required for ectopic bone and cartilage formation, LECT-1 alone induced dose-dependent muscle dedifferentiation, and their combination tuned the balance of ectopic tissues: linearly increasing cartilage with LECT-1 dose and maximizing controlled, cortical-like bone at an intermediate dose. Together, these results indicate that skeletal muscle can act as an *in vivo* incubator in which resident progenitors are guided through an endochondral-like program to generate bone and cartilage. Future studies will reduce the BMP-2 dose to favor an exclusive cartilage response and will evaluate the resulting ectopic tissues for the repair of critical-size bone defects.

## Acknowledgements

The authors thank the Bioengineering Materials Laboratory at Hofstra University for technical support. Special thanks to Jillian Haller, Nicole Newberger, Hazel Consunji de Guzman, Phoebe Christake, Evan Carroll, Andrew Tarabokija, Henna Chaudhry, Michelle Paszek, Brittany Klub, Michelle Sirochinsky, Abhiram Podili, Daniela Rios, Jodee Calder, Esther Thomas, Jessica Barayuga, and Naatram Jotis for their assistance with animal care, imaging, and histological analysis. The authors also thank Dr. Paul Vaska and his staff at Stony Brook University for the µ-CT use.

## Author Disclosure Statement

BMP-2 (Infuse) was provided by Medtronic through the External Research Program material award. R.C. de Guzman is a named inventor on a pending patent application, filed by and assigned to Hofstra University, covering methods and compositions for the use of BMP-2 and LECT-1 in cartilage and bone regeneration. The keratin biomaterial used in this study differs from the residual-hair biomaterial under separate commercial development. No other authors have any competing financial interests.

## Funding Information

This work was supported in part by the Medtronic External Research Program.

## Notes

### Competing Interest Statement

R.C. de Guzman is a named inventor on a pending patent application, assigned to Hofstra University, covering methods and compositions for the use of BMP-2 and LECT-1 in cartilage and bone regeneration. BMP-2 (Infuse) used in this study was provided by Medtronic through the External Research Program material award. The authors declare that no additional payments or services from third parties were received in the past 36 months that could be perceived to influence the submitted work. No other authors have competing financial interests.

